# Monte Carlo simulations to propagate the uncertainty of automated classification into ecological models

**DOI:** 10.64898/2026.01.14.699344

**Authors:** Célian Monchy, Olivier Gimenez, Céline Le Bohec, Gaël Bardon, Marie-Pierre Etienne

**Affiliations:** CEFE, Université de Montpellier, CNRS, EPHE, IRD, Montpellier, France; Ensai, CREST-UMR 9194, CNRS, Université de Rennes, 35000 Rennes, France; Université de Strasbourg, CNRS, IPHC UMR 7178, F-67000 Strasbourg, France; Centre Scientifique de Monaco, Département de Biologie Polaire, Monaco, Principality of Monaco

**Keywords:** Deep Learning, uncertainty propagation, Monte Carlo simulation, automatic classification, ecological inference, camera traps

## Abstract

Deep learning (DL) is increasingly integrated into quantitative ecology, particularly for automating the classification of sensor data in biodiversity monitoring. In addition to substantially reducing data processing effort, DL models often achieve high classification performance. However, despite ongoing improvements, certain species or classification tasks remain challenging, and predictions are rarely error-free. Manual verification is frequently included into data processing pipelines to mitigate misclassifications, but this approach may mask rather than quantify uncertainty.

Here, we propose directly incorporating classification uncertainty into ecological inference, rather than filtering it afterwards. Specifically, we treat model predictions as probabilistic outputs rather than fixed class assignments, by using Monte Carlo simulations to propagate uncertainty from the classification process into downstream ecological models.

We illustrate this approach using two case studies. The first estimates stochastic population growth rates for a penguin population using detection time series derived from Radio Frequency IDentification (RFID). The second propagates uncertainty in species identification from camera trap images into occupancy estimates. For both, we compare results obtained propagating classification uncertainty with those from conventional single-class attribution at two confidence score thresholds. Our findings show that propagating uncertainty typically leads to higher, more optimistic ecological estimates compared to the single-class confidence approach. Importantly, this method expands the total uncertainty interval by explicitly introducing the confidence score produced by automatic classification as a representation of uncertainty. Quantifying this uncertainty in parameter estimation allows for more informed and reliable ecological interpretations. Monte Carlo simulations offer a flexible and accessible means to integrate classification uncertainty into diverse ecological modelling workflows.

## 1 Introduction

The use of sensors and others sophisticated and high-performance instrumentation has become widespread in biodiversity monitoring, particularly for assessing ecosystems’ responses to stressors and global changes. Sensor technologies, including ground-based, remote, in situ, and wireless sensors, have advanced significantly in recent years, offering finer spatial and temporal resolution that is particularly well suited for biodiversity tracking (Marvin et al., 2016). For ecologists, sensor technologies serve as both an alternative and a complement to traditional field observations, offering new opportunities to study population dynamics, physiology, behaviour and social interactions (Wilmers et al., 2015). Among the most effective tools for wildlife passive monitoring are camera traps and acoustic recording units. When deployed as part of a sensor network, these tools enable large-scale surveys of inaccessible areas. They are non-invasive, minimizing disturbance to animal populations, and their discretion reduces biases associated with human presence (Steenweg et al., 2017). Like camera traps, acoustic recording units act as autonomous sensors. They are particularly useful for detecting species that are otherwise difficult to observe, such as small organisms that produce acoustic signals (Gibb et al., 2019). Additionally, wireless sensor networks have become essential for ecological studies, facilitating the collection of high-frequency data across a wide range of environmental parameters (Porter et al., 2005).

Sensor networks produce a wealth of data from which relevant ecological information need to be extracted. Processing sensor data to track species, individuals, or activities requires classification of recorded events. While this classification can be performed manually by experts or collaboratively through citizen science platforms (Willi et al., 2019), the ever-increasing volume of sensor data makes manual approaches increasingly unfeasible, both due to the time required and the potential for biases introduced by multiple people processing data over extended periods. As a result, automated classification using deep learning algorithms has become an essential tool. The use of artificial intelligence to classify ecological data primarily relies on neural networks (NNs), which are trained to extract features and assign them to specific categories (Norouzzadeh et al., 2018; Pollock et al., 2025). Techniques such as transfer learning and data augmentation enhance the performance of NNs in recognizing patterns within images, videos, or acoustic recordings (Duggan et al., 2021). One of the most widely used metric for evaluating NNs’ classification performance is accuracy, as it measures discrepancies between predicted and actual labels. Notably, Deep Learning (DL) models achieve high accuracy—often exceeding 90%—in species identification (Swanson et al., 2015; Norouzzadeh et al., 2018), demonstrating their broad applicability across taxa, including terrestrial and marine mammals (Schneider et al., 2024), insects (Høye et al., 2021), and birds (Ferreira et al., 2020). However, despite their high accuracy on large datasets, NNs may struggle with specific classification tasks or cryptic species. For instance, Duggan et al., 2021 reported low accuracy in identifying gray squirrels despite their high representation in the training dataset, likely due to their small size and similarity to the background. In the same study, the DL model used encountered difficulties distinguishing between visually similar species, such as dogs and coyotes, and showed variability in identifying humans, a highly diverse class. Similarly, Tabak et al., 2020, in their work on wildlife classification, observed low ability to positively identify small, elusive taxa like Passeriformes and Mustelidae. Beyond species identification, individual recognition tends to perform less reliably due to the smaller size of training datasets (e.g. Patton et al., 2023).

Although DL models achieve high accuracy in many classification tasks, errors may persist.

This is particularly concerning because classification precedes ecological analyses and can introduce biases into final inferences. Even processes that include manual verification are not completely exempt from misclassification issues. Indeed, in manual labelling, human experts are expected to flag ambiguous records, which are generally discarded from further analysis. While this reduces the risk of misinterpretation of ecological results, it also leads to data loss. Fully automated labelling, by contrast, is more prone to misclassifying ambiguous records. However, since each prediction is accompanied by a continuous confidence score, these ambiguous records can be identified - provided that some precautions are taken, like incorporating a calibration procedure to be able to interpret these scores as a risk of error (Dussert et al., 2024). Two strategies then emerge: one can, as in manual labelling, remove ambiguous records with scores below a given threshold, at the cost of further data loss (e.g. Whytock et al., 2021; Cole et al., 2022; Lonsinger et al., 2023); or one can integrate the confidence scores as measures of classification uncertainty within the labelling process (e.g. Rhinehart et al., 2022; Katsis et al., 2025; Ogawa et al., 2025). Thus, misclassification is not only related to automated data labelling, but deep learning makes the classification uncertainty explicit and also offers the opportunity to formally propagate it through downstream ecological analyses.

Many studies rely on sensor data to estimate key ecological parameters such as habitat use (e.g. Bowkett et al., 2007; Rovero & Marshall, 2009), population structure (e.g. Silveira et al., 2003), community diversity, and species abundance (e.g. Karanth et al., 2006). Reducing manual verification in automated classification saves time but raises concerns about its impact on the reliability of quantitative analyses. This trade-off between processing efficiency and inference reliability necessitates an assessment of the risks posed by classification errors. The integration of deep learning techniques into ecological monitoring has received considerable attention, particularly for tasks involving detection and classification (Cowans et al., 2024). While numerous statistical and computational methods have been explored to mitigate uncertainty in such applications (e.g., Ferreira et al., 2020; Adjei et al., 2022), approaches that use continuous confidence scores remain relatively rare.

Recent advancements have introduced models that explicitly utilize continuous confidence scores, especially in acoustic monitoring contexts. For instance, Balantic and Donovan, 2019 proposed to aggregate confidence scores at the survey level, applying thresholds to these aggregates to estimate the probability of true species identification. Similarly, Singer et al., 2024 examined the precision of various thresholding techniques applied to scores generated by a free audio bird classifier, and modelled optimal thresholds for each target species, to minimize false positive detections.

In the field of occupancy modelling, Cole et al., 2022 compared the outputs of different model designs using BirdNET versus manually annotated data. Their results underlined the benefits of filtering automatic annotations with species-specific confidence thresholds to reduce false positive detections. They also integrated confidence score distributions, as suggested by Kery and Royle, 2020, to inform the probability that a detection labelled as positive by BirdNET is truly positive.

Further extending this approach, both Rhinehart et al., 2022 and Kery and Royle, 2020 developed occupancy models that treat continuous confidence scores from automated classifiers as Gaussian mixtures. Rhinehart et al., 2022 integrated these scores to determine the likelihood that a given file contains the target species and proposed enhancing model performance by including a subset of manually annotated detections for each site.

These approaches have the advantage of incorporating information about the label prediction confidence and explicitly propagating it into the final inference. However, such continuous score-based methods are primarily used in species identification within the site occupancy modelling framework and often require structural modifications to the ecological model. In the perspective of informing ecological inferences with uncertainty about the automatic classification process, we propose a Monte Carlo simulation-based method to propagate classification uncertainty into model estimates. Like other bootstrap methods, Monte Carlo simulations rely on sampling from specified distributions to approximate input data variability. After a brief overview about DL models for classification, we explain the principles behind the Monte Carlo method. Through two case studies, we demonstrate how this flexible approach quantifies the impact of classification uncertainty on ecological parameter estimation.

## 2 Method : the propagation of classification uncertainty with Monte Carlo simulations

### 2.1 Neural Networks

Neural Networks (NNs) can be thought of as highly complex models where inputs are processed through multiple layers of neurons in a latent space. Automatic classification with DL models is inherently deterministic, once trained, parameter values are fixed (though typically opaque to the user), ensuring that classifying identical input data results in the same predictions. In multi-class classification tasks, the NN’s output is a loading vector *z* = (*z*_1_, …, *z*_*K*_) whose dimension equals the number of classes, this vector is classically transformed through a softmax function *σ* into confidence scores of classification *σ*(*z*_*i*_) for each of the *K* class. The ability of this score to reflect the predictive uncertainty of the neural network (NN) depends on its calibration, which is related to the model’s architecture and the amount of training data, for example (Minderer et al., 2021). Based on the hypothesis that confidence scores reflect calibrated probabilities, we employed Monte Carlo (MC) simulations to reflect the uncertainty at the classification output level and propagated this uncertainty through statistical models to evaluate its impact on ecological inferences.

### 2.2 Monte Carlo simulations

Uncertainty propagation using the MC method relies on random sampling techniques to account for variability in observation data due to automatic classification uncertainty. Class assignment follows a Bernoulli distribution, where the calibrated confidence score represents the probability of success. For multi-class classification questions, such as species recognition, the classification task is transformed into multiple binary classifications, focusing on each class of interest in a *one-vs-all* strategy. By sampling class assignments based on scores rather than selecting the most credible class, we introduce variability into the dataset. Using the diversity of the so produced dataset, we are able to propagate the initial uncertainty of data classification into the outputs of the ecological models.

To illustrate MC method for propagating uncertainties from classified data into ecological model estimates, we compared the probabilistic interpretation of the confidence score with a single-class confidence approach involving the application of a threshold to the confidence score. We examined the influence of two thresholds on the estimates: 0.5 and 0.95. A threshold of 0.95 corresponds to filtering predictions for which the classifier is highly confident, whereas a threshold of 0.5 corresponds to selecting the most likely class, regardless of the classifier’s confidence. Finally, we investigated the relative contribution of classification and parameter estimation uncertainty to the total uncertainty. The model uncertainty was quantified by the confidence interval, recorded for each estimate from MC simulation and the single-class confidence approach. The total uncertainty was determined as the range between the minimum and maximum bounds of all confidence intervals (CI), including the results from both interpretation of the confidence score.

## 3 Case studies

To illustrate the flexibility of the MC approach, we developed two case studies. Firstly, we propagated the uncertainty related to behaviour pattern identification into the growth rate of a population of seabirds as calculated in a population matrix model (Bardon, 2024). In a second case study, we addressed the uncertainties associated with automated species classification in camera trap images, and their propagation into a site occupancy model (Gimenez et al., 2022).

### 3.1 Demographic growth rate from time series of RFID detections

We focus on a population of king penguins (*Aptenodytes patagonicus*) located in the Crozet archipelago (Terres Australes et Antarctiques Françaises) and long-term monitored within the framework of the IPEV 137 and CNRS SEE-Life programmes. Individual movements are tracked with passive integrated transponder (PIT) tags (Le Bohec et al., 2008; Bardon et al., 2023). Radio-Frequency IDentification (RFID) technology records every transit of tagged birds between the sea and their breeding colony, generating lifelong time series. Given the highly stereotyped reproductive movement patterns of these seabirds, individual breeding characteristics can be inferred from detection time series (Descamps et al., 2002). Bardon et al., 2023 developed RFIDeep, a workflow based on automatic classification that extracts movement patterns from RFID data and classifies individuals based on their breeding activity. Two independent convolutional neural networks determine both the breeding status (breeder / non-breeder) and breeding outcome (failure / success). A demographic matrix model provides an age-structured representation of the population with fluctuating vital rates (fecundity and survival) and admits short-term and long-term forecasting of population size thanks to the estimation of the population growth rate (Cohen, 1986). We focused on the stochastic growth rate approximation to evaluate long-run population growth in light of the influence of a fluctuating environment on vital rates (Tuljapurkar et al., 2003; Caswell, 2006).

#### 3.1.1 Population matrix modelling

RFID data have been collected from tagged individuals since 1998 and have been used to estimate population parameters such as survival and fecundity (Le Bohec et al., 2008). Here, we focused solely on individuals tagged as chicks (N=11877), employing an age-classified pre-breeding census matrix model (Caswell, 2006). Assumptions for the model included that an individual was considered dead if no signal was detected for two consecutive years and demographic rates remained constant after the age of 10. For each year, the survival rate *s* = (*s*1, …, *s*10+) was defined as the proportion of individuals that survive the following year, while fecundity *f* = (*f* 1, …, *f* 10+) was measured as the proportion of individuals classified as having achieved breeding success. For each year from 2008 to 2021, we built the transition matrix *A*(*t*), defined as *N* (*t*+1) = *A*(*t*)*N* (*t*), with *N* (*t*) the composition of the population at time *t*.

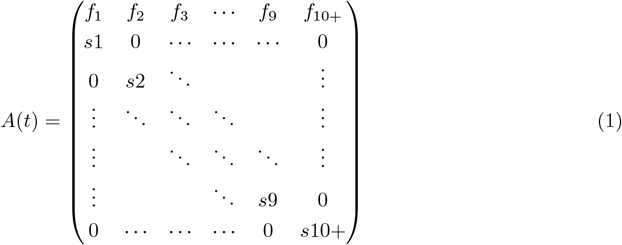

We then used the *popbio* R package (Stubben & Milligan, 2007) to approximate the stochastic growth rate and its 95% CI (Tuljapurkar & Haridas, 2006), which represented the time-averaged cumulative growth of the population in a temporally stochastic environment (Tuljapurkar et al., 2003). The 95% confidence interval was derived from standard errors computed with bootstrap, to account for sampling variance due to variations in vital rates (survival and fecundity)(Hendi, 2023; Hernandez-Suarez & Rabinovich, 2024).

#### 3.1.2 Propagation of uncertainties in breeding activity classification and quantification of the total uncertainty in stochastic growth rate

In this population model, fecundity was defined as the proportion of individuals contributing to reproduction. The contribution of each individual was determined by a sequence of two binary classifications - breeding status and breeding outcome - based on detection time series. Uncertainties associated with automatic classifications were accounted for and propagated in the calculation of fecundity using the probabilistic approach of MC. For an individual, the reproductive success means being a breeder and ensuring chick survival during the breeding cycle period of more than one year long. Therefore we sampled the breeding outcome of each individual as a Bernoulli random variable, where the probability of success was the product of the confidence scores for both classification tasks. We ran 100 MC simulations, drawing the breeding outcome for each individual and we compared the resulting distribution of mean fecundity per year with the results of the single-class confidence approach for two threshold values (0.5 and 0.95). Finally, we used the fecundity values resulting from the MC simulations to generate 100 sets of transition matrices and computed the stochastic growth rate for each of them. Similarly, we calculated the stochastic growth rate from transition matrices produced using the single-class confidence approach to breeding activity.

### 3.2 Occupancy modelling with pictures from camera traps

We focused on a reintroduced population of Eurasian lynx (*Lynx lynx*) currently recolonizing the French Jura Mountains. The monitoring survey was conducted by the *French Office for Biodiversity (OFB)*, in collaboration with the *Departments of Hunting Federations from Jura, Ain, and Haute-Savoie*. Camera traps were deployed at 29 monitoring sites across the Jura Mountains, each site being equipped with two cameras to maximize detection probability. Data collection spanned the entirety of 2017 and continued thereafter, ensuring a monitoring period of at least one year. In total, 68,402 images were manually annotated before being classified using the automated recognition tool *DeepFaune* (Rigoudy et al., 2023).

#### 3.2.1 Single-species occupancy modelling

To estimate the proportion of area occupied by the Eurasian lynx controlling for its imperfect detection, we used a single-species occupancy model (MacKenzie et al., 2002). The occupied area is defined as the proportion of occupied sites. In order to challenge our method on contrasted dataset, we considered two additional taxa: chamois (*Rupicapra rupicapra*) and lagomorphs represented by the European hare (*Lepus europaeus*). The detection of a species at site *i* during the sampling occasion *j*, denoted as *Y*_*ij*_, was defined conditionally on the latent occupancy state *Z*_*i*_ of site *i* :

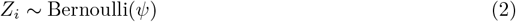

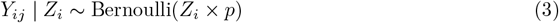

where *p* and *ψ* represented the detection and occupancy probabilities, respectively, which were assumed constant throughout the year and across sites. Data were aggregated by month, such as the target species was considered detected (*Y*_*ij*_ = 1) when it is identified on at least one picture *k* during the *j*^*th*^ month, at site *i* :

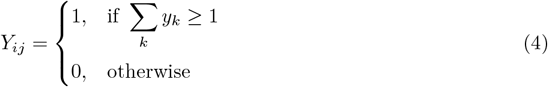

We used the R package `unmarked` (Fiske & Chandler, 2011) to implement the occupancy model within the maximum likelihood framework.

#### 3.2.2 Propagation of uncertainties in species recognition and quantification of the total uncertainty in occupancy parameters

We examined the influence of confidence score interpretation on occupancy estimates. In the probabilistic approach used to propagate species recognition uncertainties, we tested multiple aggregation methods to accommodate with identification and detection levels (see Supplementary Methods). We finally kept the most intuitive in which the identification of the target species on a picture follows a Bernoulli random variable with the confidence score associated with the species’ class as the probability. We conducted 100 MC simulations, recording for each simulation the detection history per site, the maximum likelihood estimates of both detection and occupancy parameters, and the derived 95% CIs from the Hessian matrix. In the single-class confidence approach involving the application of a threshold value (0.5 and 0.95), the target species was considered present if its confidence score exceeded the threshold and absent otherwise. Finally, combining both probabilistic and single-class confidence approach, we investigated the relative proportion of classification and parameter estimation uncertainties to the total uncertainty.

## 4 Results

### 4.1 Demographic growth rate from time series of RFID detections

Annual average fecundity estimates derived from the single-class confidence approach were consistently lower, particularly under more restrictive threshold values (Figure 1). Discrepancies are observed between the two approaches since the estimates resulting from MC simulations are systematically above the estimates from the single-class confidence method. For example, using a 0.5 confidence threshold in the single-class method produced fecundity values in average 13% behind the mean value obtained from MC simulations. When the threshold was increased to 0.95, the single-class estimate fell on average 15% below the MC-derived mean. MC simulations also provided a quantification of uncertainty associated with both annual breeding rate and breeding success, resulting in relatively high confidence ranges for average fecundity across years. This uncertainty spanned from 40% of total fecundity in 2018 to 13% in 2019, indicating a non-negligible impact of uncertainty propagation on long-term fecundity trends.

**Figure 1:**
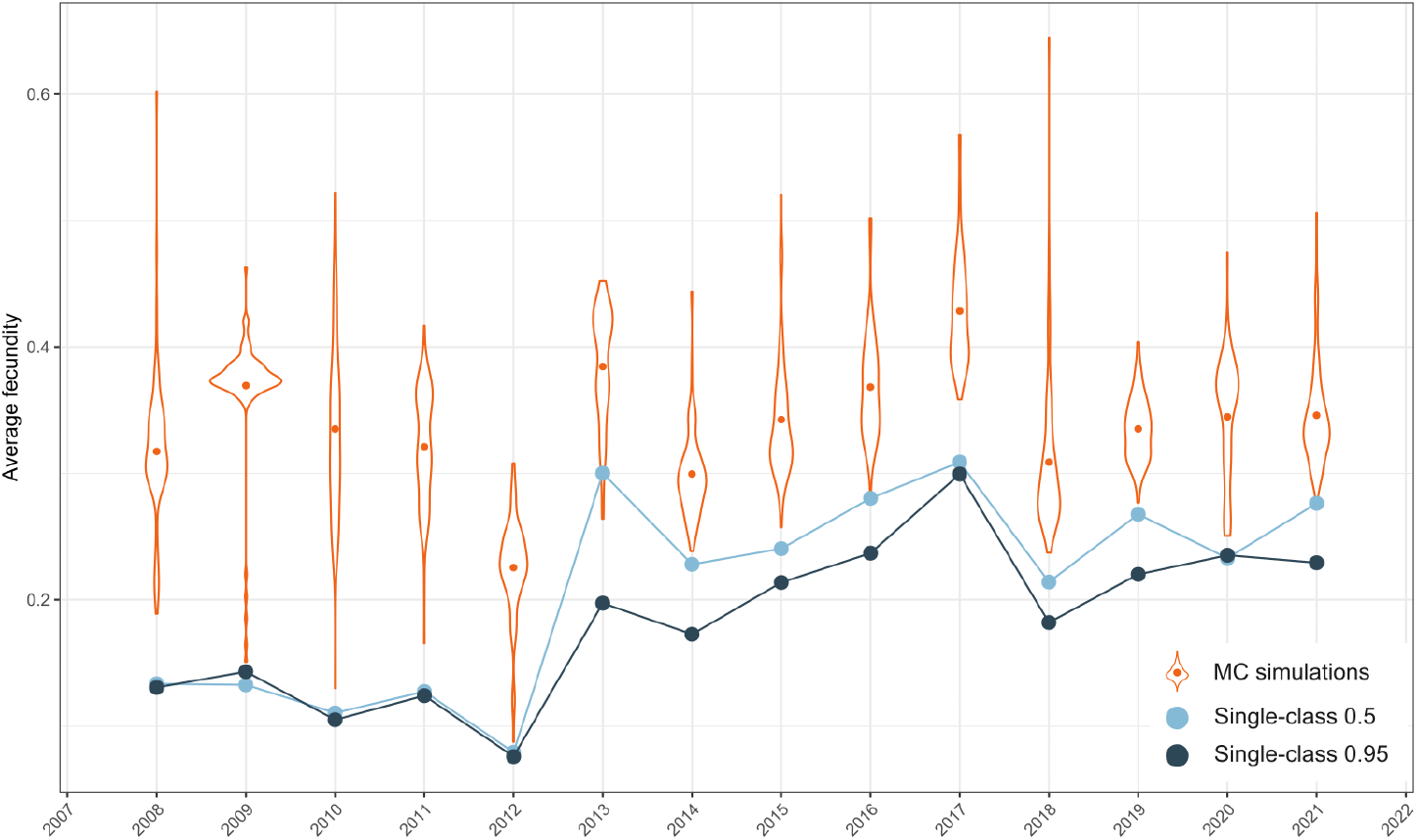
Annual variation in the average fecundity of a king penguin population. Fecundity was calculated for each year as the proportion of individuals that successfully reproduced relative to the number of individuals breeding that year. Breeding success for each individual was determined through an automatic classification of movement patterns during the breeding period. Classification uncertainty regarding breeding outcomes was addressed using MC simulations (orange violin plots, dots represent distribution’s mean). The single-class confidence approach did not consider the classification uncertainty of the breeding outcome, and it is based on two threshold values : 0.5 (light blue dots) and 0.95 (dark blue dots).

The propagation of fecundity uncertainty had a significant impact on the approximation of the stochastic growth rate of the population. In contrast, the confidence interval reflecting the sampling variation in vital rates used to approximate the stochastic growth rate from a single set of transition matrices is negligible. Applying the single-class confidence approaches, the stochastic growth rates were estimated at 6.5% and 5.1% for threshold set at 0.5 and 0.95 respectively (Figure 2). When uncertainties were propagated with MC simulations the stochastic growth rate fluctuated between 9.5% and 29.0%.

**Figure 2:**
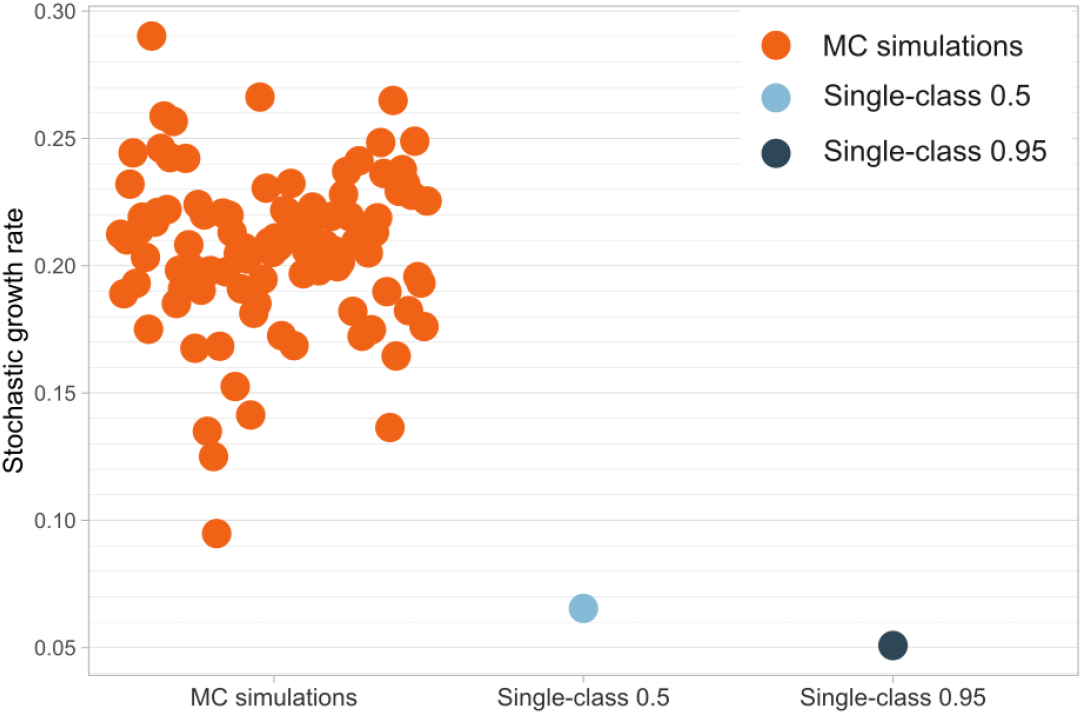
Tuljapukar’s approximations of the stochastic growth rate of a king penguin population. The stochastic growth rate was approximated using population matrices from 2008 to 2021. The probabilistic approach, based on 100 MC simulations, accounted for uncertainty about breeding outcome classification (left). The single-class confidence approach was based on two threshold values : 0.5 (middle) and 0.95 (right).

### 4.2 Occupancy modelling with pictures from camera traps

Detection estimates fluctuated continuously across MC simulations, while occupancy estimates took discrete values due to the variation in the number of sites classified as occupied from one simulation to another. Lynx occupancy was characterized by three distinct modes, corresponding to the occupation of 25, 26, or 27 out of 29 sites (Figure 3). In contrast, occupancy estimates for the other two species displayed more subtle multimodal distributions, with more than four distinguishable peaks. Some sites were classified as occupied in only a small number of simulations due to isolated identifications (see Supplementary Results). These rare detections influenced the overall occupancy estimates by causing shifts to higher occupancy values.

**Figure 3:**
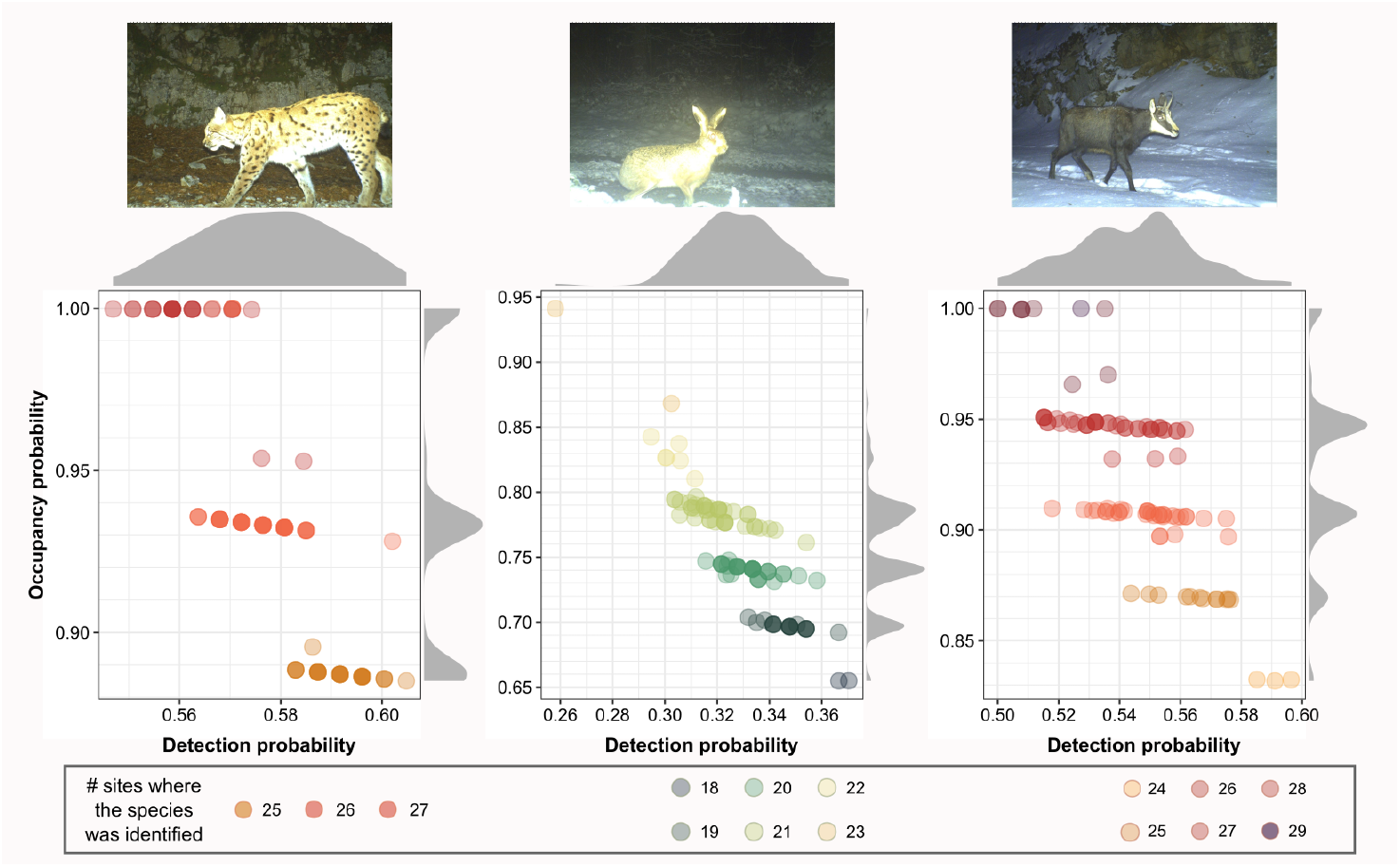
Occupancy and detection estimates from 100 MC simulations for lynx, lagomorph and chamois. The parameters of the occupancy model were estimated in 100 MC simulations (points) in which the scores were randomly drawn for each pictures. In each simulation, the number of occupied sites was determined according to the identifications which varied from simulation to simulation. Density distributions for both occupancy (right margin) and detection (top margin) estimates are based on every MC simulation.

Detection estimates were equivalent for the two approaches, regardless of the confidence score threshold, and similarly for all the three species. For lynx and lagomorph, occupancy estimates from the single-class confidence approach consistently fell within the range obtained via MC simulations. In contrast, raising the confidence score threshold in the single-class confidence approach reduced occupancy estimates for both lagomorph and chamois (Figure 4).

**Figure 4:**
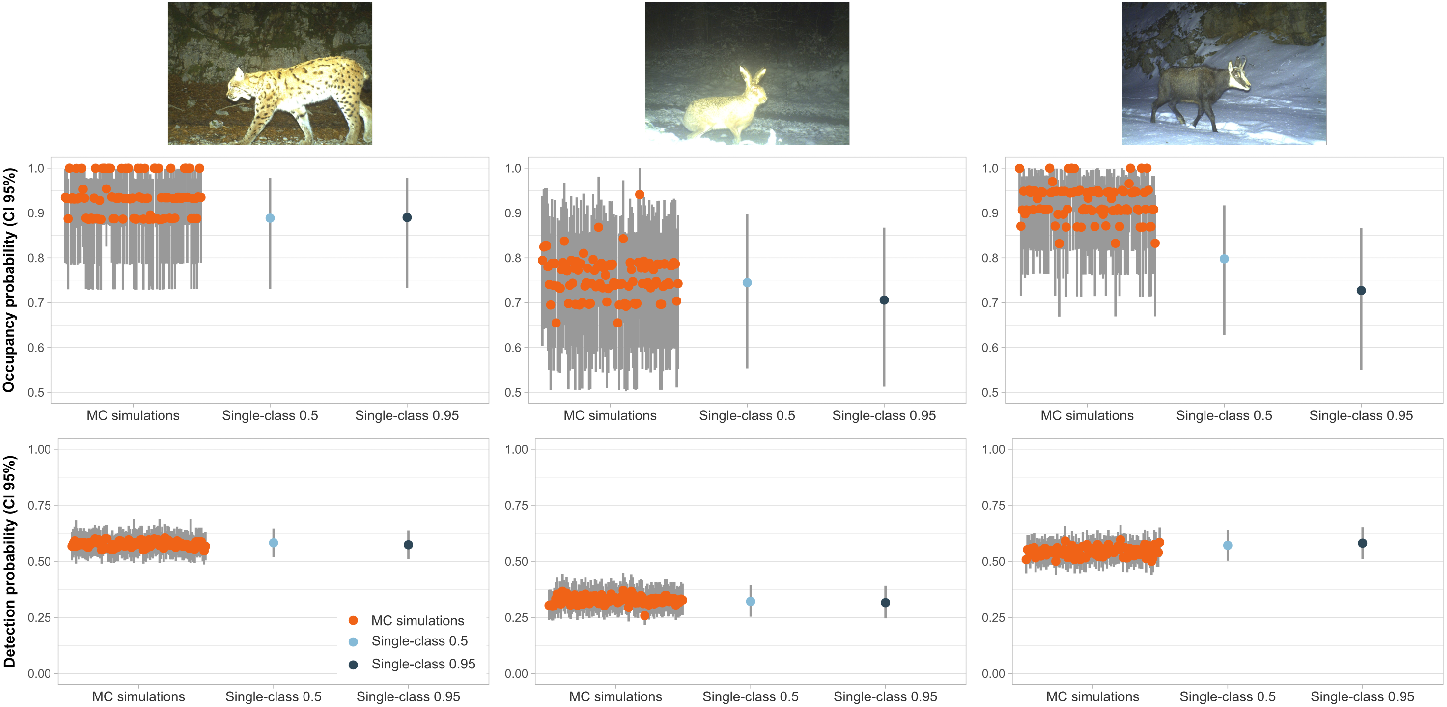
Estimates of occupancy and detection probabilities for lynx, lagomorph and chamois. The single-class confidence approach was based on two threshold values : 0.05 and 0.95. The probabilistic approach, based on 100 MC simulations, accounted for uncertainty about automatic species recognition on images. Vertical segments represent the profile likelihood 95% CIs computed for occupancy and detection parameters.

**Figure 5:**
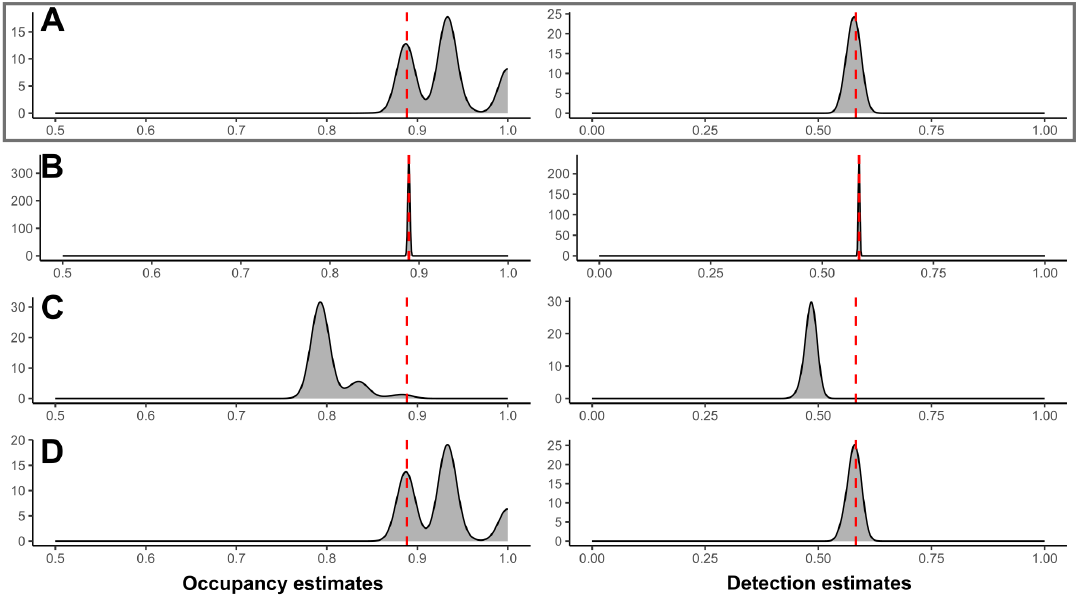
Comparison of aggregation methods for score use at occasion level based on estimates using the probabilistic approach of occupancy and detection probabilities for lynx. In **A** and **B**, a lynx was considered detected if it was identified at least once during the occasion. The identification on an image was drawn in a Bernoulli distribution where the parameter was **(A)** the score, or **(B)** the score weighted by the likelihood that the site was occupied. In order to take into account the detection history to give prior information about the identification, in **C** the identification on an image was weighted by the probability the site was occupied given the detection history where the current image is removed. Finally, in **D** the detection state for each occasion was estimated according to the method proposed by Balantic and Donovan, 2019, where detection is defined by the probability the species has been identified at least once during the occasion, considering the scores of images from the occasion. The aggregation methods are compared with the single-class confidence estimates (red dashed line). The method used in the probabilistic approach of the paper is **A** (grey frame).

**Figure 6:**
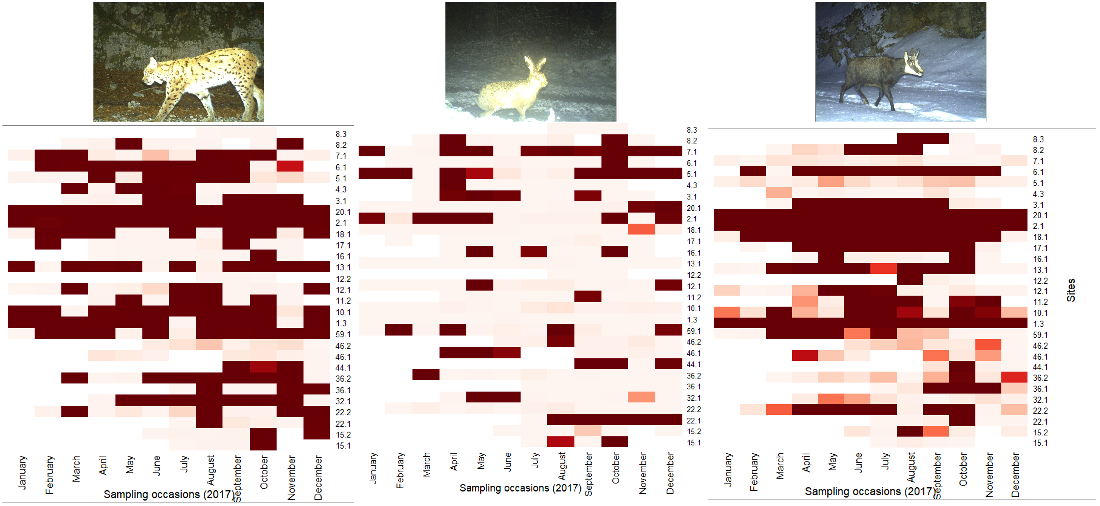
Matrix of identifications resulting from 100 MC simulations for lynx, lagomorph and chamois. The number of identifications was qualified per sites (y-axis) and sampling occasions (x-axis). Darker cells determine a high number of simulations in which the species was identified, when white cells determine that no data was available for this occasion of this site.

Many images of chamois were assigned a confidence score below 0.95, indicating an ambiguous classification. As a result, the MC simulations produced higher occupancy probability estimates than those derived from the single-class confidence approach, especially when high thresholds were applied. Although detection estimates for chamois remained largely unchanged, occupancy probability was estimated at approximately 0.8 [0.63; 0.92] with a 0.5 threshold, with only 19% of the MC-derived estimates falling within this 95% CI. For chamois, this proportion decreased further at 3% with a 0.95 threshold, though the overall difference between the approaches was not statistically significant. The degree of overlap between classification uncertainty and model uncertainty varied by species and parameter. For lynx detection probability, the model uncertainty defined by the 95% CI encompassed all estimates derived from the MC simulations. Here, the aggregate uncertainty ranged from 0.49 to 0.69, with model uncertainty contributing approximately 57% of this range (Figure 4). Comparable trends were noted for lagomorph and chamois detection estimates (ranges from 0.22 to 0.44 and from 0.44 to 0.66, respectively), with model uncertainty representing 56% and 57% of the corresponding detection ranges.

For lynx and lagomorph, the model uncertainty determined using the single-class confidence approach aligned well with the ranges of classification uncertainty derived from the MC simulations. However, for chamois, the classification uncertainties from MC simulations did not overlap with the 95% CIs from the single-class confidence estimates, indicating that model uncertainty contributed to increase the total uncertainty due to the discrepancies between the approaches.

## Discussion

We used Monte Carlo simulations to quantify the impact of automatic classification uncertainties on ecological inferences. When applied to the confidence scores from classification, this method is particularly valuable, as it integrates the variability of input data-resulting from automatic classification-into the estimation of ecological parameters.

We defined reference points as the estimates obtained from the dataset, where the labels are determined by thresholding the classification score. We compared those reference points with results from the MC simulations. The propagation of uncertainty through MC simulations and the single-class confidence approach result from two distinct interpretations of the confidence score. We considered both approaches when defining total uncertainty for two ecologically meaningful parameters: the probability of presence and the stochastic population growth rate. We initially expected the probability density function of the estimates obtained via simulations to be centred around the reference estimate derived from the single-class confidence approach. However, we found that MC simulations led to higher estimates of ecological parameters compared to the approach without propagating classification uncertainties.

In the first case study, fecundity was linked to the number of successful breeders, itself derived from two automatic classifications: breeding status and breeding success. Propagating classification uncertainties for both breeders and successful breeders brings more variability to the calculation of annual fecundity than the single-class confidence approach. In this context, not propagating classification uncertainties might be considered as a conservative approach for estimating the stochastic growth rate of a threatened population. Slight underestimation of growth could favour more cautious conservation strategies. Conversely, underestimating the growth of a harvested or invasive species may lead to suboptimal management decisions.

Propagating uncertainty on species recognition led to higher estimates for the occupancy probability, across all studied species. This finding highlighted the utility of probabilistic methods to account for classification errors in hierarchical models not explicitly structured to handle them. While false negatives were more common, false positives could still occur, even in detection data collected via traditional field methods. The effects of false positives on occupancy estimates have been discussed in depth (McClintock et al., 2010; Miller et al., 2011), and various methods have been proposed to account for them, including explicit modelling (e.g, Royle and Link, 2006) and adapted study design (e.g., Chambert et al., 2015). More recently, a growing body of work (e.g., Guillera-Arroita et al., 2017; Wright et al., 2020) has emphasized treating the identification process as distinct from detection when modelling occupancy. For occupancy studies based on sensor data classified with deep learning, another approach involves staying with the traditional framework of occupancy models (MacKenzie et al., 2002) while reducing misclassification risks. For instance, Whytock et al., 2021; Lonsinger et al., 2023 demonstrated that applying high confidence thresholds to scores reduced false positives, leading to more reliable occupancy estimates. However, Lonsinger et al., 2023 also reported an exception: for one species frequently confused with another, even high thresholds resulted in occupancy overestimation.

This highlights the importance of propagating classification uncertainty, especially for difficult tasks of classifications, like distinguishing cryptic or visually similar species. In such cases of confidence score application, it becomes essential to ensure that scores accurately reflect the likelihood of correct predictions. Poorly calibrated classifiers can produce overconfident but incorrect predictions, which can lead to biased ecological estimates. Calibration metrics such as the Brier score and Expected Calibration Error are commonly used to quantify the alignment between predicted confidence and actual correctness (Guo et al., 2017; Minderer et al., 2021). Miscalibration can arise from various factors, including model architecture, training dataset, model size, and normalisation techniques. However, identifying optimal configurations that simultaneously maximize both classification accuracy and calibration remains challenging (Guo et al., 2017). To address this, Dussert et al., 2024 proposed post-hoc calibration methods that refine predicted confidence scores without requiring the NN to be retrained. In addition to using temperature scaling, they suggested aggregating information at the sequence level in camera trap data, rather than relying on image-level outputs. Well-calibrated confidence scores provide a more robust basis for downstream ecological analyses, especially when these scores associated with the predicted class label are employed as probabilistic uncertainties. Although calibration has been widely studied in the machine learning literature (e.g. Guo et al., 2017; Minderer et al., 2021), our approach considers confidence scores as informative inputs to ecological models, regardless of whether they are perfectly calibrated.

Also, we showed that the combination of propagating classification uncertainties with MC simulations and increasing the classification threshold are divergent approaches in a way that it increases the amplitude of the total uncertainty. We demonstrated that propagating classification uncertainties via MC simulations and increasing the classification threshold had contrasting effects on the results. The former tended to yield more optimistic estimates by incorporating a broader range of predictions, whereas the latter adopted a more conservative approach by retaining only high-confidence classifications. When combined, these methods amplified the overall range of uncertainty, thereby highlighting a common feature of statistical modelling: the trade-off between accuracy and precision.

In the population dynamics case study, the differences in stochastic growth rate estimates were relatively high and biologically meaningful (Fig. 2). For comparison, Nelson et al., 2010 reported uncertainty in growth rates for U.S. red fox populations ranging from −1% to +16%. In our study, growth rate uncertainty ranged from 5% to 29%, illustrating the level of precision achievable when total uncertainty is explicitly accounted for.

In contrast, the second case study based on species identification, revealed a more pronounced impact. Combining both approaches substantially broadened the total uncertainty range. Propagating classification uncertainty through MC simulations allowed for the inclusion of ambiguous confidence scores, which increased overall occupancy estimates. Conversely, applying a restrictive threshold (e.g., 95%) reduced occupancy estimates, particularly for the chamois.

While Whytock et al., 2021 demonstrated that increasing classification thresholds enhances concordance between automated and expert labels, this does not guarantee correct predictions in the context of sensor data, even with manual verification. In addition to sources of error intrinsic to records such as quality and inconsistent labelling, certain species are consistently misclassified, and high thresholds may not sufficiently mitigate occupancy overestimation. In some contexts, considering the estimates from manually verified data as the truth might not be an evidence, as DL models can outperform human annotators in detecting discret patterns (e.g. Miller et al., 2023; Otsuka et al., 2024). Therefore, while lowering the threshold may appear to enhance precision, it does not necessarily lead to more accurate estimates.

Ultimately, neither approach alone resolves the inherent trade-off between accuracy and precision. However, their combination offers a more comprehensive understanding by explicitly expanding the bounds of total uncertainty.

It is important to recognize that the total uncertainty we estimated does not cover all potential sources. In ecological modelling, uncertainty arises from multiple factors: sampling error, measurement error, model structure, parameter calibration, and inherent stochasticity (Schuwirth et al., 2019). Our approach integrates classification error and sampling variability, which are particularly relevant for sensor-derived data. Ultimately, propagating classification uncertainties is especially valuable for tasks where automated classification is less reliable. To do so, we recommend two conditions be met: (1) confidence scores must be properly calibrated, and (2) the ecological model must not be biased toward one type of classification error.

Incorporating classification uncertainties using MC simulations is a promising way to improve the accuracy of ecological inferences. While this comes at the risk of reducing precision, it provides a more complete view of uncertainty, particularly when classification errors are likely or difficult to avoid. Although manual verification can reduce misclassifications, it is not error-free (Zett et al., 2022), and in some situations automated classifiers can even outperform human experts. Because the MC approach is light and easy to implement, it is well-suited for propagating uncertainty in these studies. Explicitly reporting the share of total uncertainty that comes from automatic classification will, therefore, support more informed decision-making.

## Author contributions

CM, OG and MPE contributes to the conceptualisation. CM performed analyses and wrote the original draft. OG and MPE validated, wrote reviews and edited the manuscript. CLB and GB provided RFID data, built the simplified population model and wrote reviews.

## Acknowledgements

CM benefits of a PhD grant from the CNRS through the MITI interdisciplinary program and the PITACA project. This research is funded by Biodiversa, the European Biodiversity Partnership, in the context of the Big_Picture project under the 2022-2023 BiodivMon joint call. It is co-funded by the European Commission (GA No. 101052342) and the French Agence Nationale de la Recherche. The long-term monitoring of the king penguin population from the Crozet archipelago was supported by the Institut Polaire Français Paul-Emile Victor (IPEV) within the framework of the project 137-ANTAVIA-POLAROBS, by the Centre National de la Recherche Scientifique (CNRS) through the Programme Zone Atelier de Recherches sur l’Environnement Antarctique et Terres Australes Subantarctique (ZATA) and the long-term Studies in Ecology and Evolution (SEE-Life), and by the Centre Scientifique de Monaco (CSM). We are deeply grateful to all the wintering and summering members of project IPEV 137 and all the other colleagues and students within the team, who participated in the long-term monitoring as part of the project IPEV 137 over the years. We also sincerely thank the IPEV logistics teams for their important and continued support in the field. This study was approved by the French Polar Environmental Committee and permits handling animals and access breeding sites were delivered by the Terres Australes et Antarctiques Françaises (TAAF). The authors thank the Federations of Hunters from the Jura and Ain counties for sharing the lynx camera-trap data. Data collection was carried out through the Lynx Predator Prey Program, which was funded by Auvergne-Rhone-Alpes Region, Ain and Jura departmental Councils, the French National Federation of Hunters, French Environmental Ministry based in Auvergne-Rhone-Alpes and Bourgogne-Franche-Comte Regions and the French Office for Biodiversity. The authors sincerely thank Gaspard Dussert for provided confidence scores from DeepFaune for camera-trap data. The authors are very grateful to Gaël Bardon and Céline Le Bohec for having shared their data about modelling and for having enriched our case study with reflections on the population dynamics model.

## Supplementary Methods

We tested different methods for using scores defined at the image level in the probabilistic approach of the occupancy model, where the detection state is a binary variable defined at the occasion level.

1. The most intuitive method is to use the score as the probability of identifying the species on a record, and define that the species was detected during an occasion if it was identified at least once. In other words if ∑*y*_*k*_ *>* 1 with *Y*_*ij*_ = (*y*_1_, …, *y*_*K*_), then the sum of the identifications for an occasion *j* was greater than 1 (Fig A).
2. We gave more weight to the complete detection history when drawing the identification in each MC simulation. Therefore, we weighted the scores for a site by the likelihood of that site being occupied, given its detection history as determined by the single-class confidence approach (Fig B). Each MC simulation is equivalent to *y*_*k*_ *∼ Bernoulli*(*score*_*k*_ *× P* (*Z*_*i*_ = 1|*Y*_*i*_)).
3. We tested another method to take into account the information in all images of an occasion. To draw the identification state of each image, we weighted the score by the probability that at least one other image gave a positive identification (Fig C). Then, for a MC simulation of the identification state for the record *k*_0_, we draw 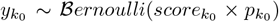 where 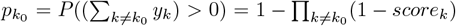 is the probability that there was at least one identification at occasion *j*, excluding *k*_0_.
4. We adapted the technique proposed by Balantic and Donovan, 2019 to not sample the species identification at the image level, but to condense the scores at occasion level by sampling the detection with the probability that the species was identified at least once during the occasion. This gave *Y*_*ij*_ *∼ Bernoulli*(1 *−* ∏_*k*_(1 *− score*_*k*_)) where ∏_*k*_ (1 *− score*_*k*_) was the probability that the species was identified on any record (Fig D).

Finally, we used the first method in the probabilistic approach, as the most intuitive and correct.

## Supplementary Results

